# Emerging role of mesenchymal stem cell-derived extracellular vesicles to ameliorate hippocampal NLRP3 inflammation induced by binge-like ethanol treatment in adolescence

**DOI:** 10.1101/2023.11.07.565776

**Authors:** Susana Mellado, María José Morillo-Bargues, Carla Perpiñá-Clérigues, Najoua Touahri, Francisco García-García, Victoria Moreno-Manzano, Consuelo Guerri, María Pascual

**Author notes:** **Corresponding author:** María Pascual, Department of Physiology, School of Medicine and Dentistry, University of Valencia, Avda. Blasco Ibáñez, 15. 46010-Valencia, Spain.

## Abstract

NOD-like receptors are innate immunity sensors that provide an early and effective response to pathogenic or injury conditions. However, abnormalities in these receptors may cause excessive inflammation. Our studies have reported that an activation of the NLRP3-inflammasome complex in ethanol-treated astrocytes and in chronic alcohol-fed mice could be associated with neuroinflammation and brain damage. Considering the therapeutic role of the molecules contained in the extracellular vesicles (EVs) derived by mesenchymal stem cells (MSC-EVs), the present study aims to evaluate whether the intravenous administration of MSC-EVs from adipose tissue, through inhibiting the NLRP3 inflammasome activation, is capable of reducing hippocampal neuroinflammation in adolescent mice treated with binge drinking. We demonstrate that MSC-EVs ameliorate the activation of the hippocampal NLRP3 inflammasome complex and other NLRs inflammasomes (e.g., NLRP1, NLRC4 and AIM2), as well as the alterations of inflammatory genes (IL-1β, IL-18, iNOS, NF-κB, MCP-1 and CX3CL1) and miRNAs (miR-21a-5p, miR-146a-5p and miR-141-5p) induced by binge-like ethanol treatment in adolescent mice. Bioinformatic analysis further revealed the involvement of miR-21a-5p and miR-146a-5p with inflammatory target genes and NOD-like receptor signaling pathways. Taken together, these findings provide, for the first time, evidence of the therapeutic potential of MSC-derived EVs to restore the hippocampal neuroinflammatory response through the NLRP3 inflammasome activation induced by binge drinking in adolescence.

## INTRODUCTION

Nucleotide-binding oligomerization domain (NOD)-like receptors (NLRs) are one of the major forms of innate immune sensors that provide immediate responses against pathogenic invasion, tissue injury and stress conditions. The NLRs family is activated through the recognition of both pathogen-associated molecular patterns (PAMPs) and damage-associated molecular patterns (DAMPs) in the host cytosol (Saïd-Sadier and Ojcius, 2012). Several members in the NLRs family have been identified, such as NLRP1 (pyrin domain-containing 1), NLRP3, NLRC4 (caspase recruitment domain-containing 4), and AIM2 (a member of the PYHIN protein family). Activation of NLRs are capable of forming inflammasomes or multiprotein complexes that activate caspase-1, leading to the processing and secretion of pro-inflammatory cytokines interleukin-1β (IL-1β) and IL-18 (Almeida-da-Silva et al., 2023).

Among NLRs, NLRP3 is currently the most fully described inflammasome. It consists of the NLRP3 scaffold, adaptor ASC (apoptosis-associated speck-like protein containing a CARD) and caspase-1 (Almeida-da-Silva et al., 2023). The activation of these receptors triggers an immune response activation and tissue repair processing. However, abnormalities in any of these inflammasome receptors may cause excessive inflammation due to either hyper-responsive innate immune signaling or sustained compensatory adaptive immune activation (Wang and Hauenstein, 2020; Piancone et al., 2021). Our previous studies demonstrated that ethanol activates the NLRP3-inflammasome complex, in both cultured astroglial cells and in cerebral cortices of WT (wild-type) mice, triggering caspase-1 activation and the induction of IL1β and IL-18, which leads to neuroinflammation and brain damage (Alfonso-Loeches et al., 2014, 2016).

One of the most important brain areas to develop in adolescence is the hippocampus, which is extremely vulnerable to the effects of binge ethanol drinking in both humans (Nagel et al., 2005; Quigley and Committee on substance use and prevention, 2019) and rodent adolescents (McClain et al., 2011; Mira et al., 2019). Our previous studies have demonstrated that binge-like ethanol exposure in adolescence, through the activation of innate immune receptors TLR4 (Toll-like receptor 4) in glial cells, can lead to the release of cytokines and chemokines, causing neuroinflammation and neural damage (Fernandez-Lizarbe et al., 2009; Alfonso-Loeches et al., 2010). Activation of the TLR4 response has also been associated with memory and learning dysfunction induced by binge drinking in adolescent mice (Montesinos et al., 2015). Notably, our findings have demonstrated that the extracellular vesicles (EVs) derived by mesenchymal stem cells (MSC-EVs) restore the neuroinflammatory response, along with myelin and synaptic structural alterations in the prefrontal cortex, as well as cognitive and memory dysfunctions induced by binge-like ethanol treatment in adolescent mice (Mellado et al., 2023).

Therefore, by considering the therapeutic role of the molecules contained in the MSC-EVs, which participate in physiological and pathological conditions (e.g. neurodegenerative diseases) (Yin et al., 2019), the present study provides evidence that the intravenous administration of MSC-EVs from adipose tissue ameliorates the NLRP3 inflammasome-mediated neuroinflammation and inflammatory-related genes and microRNAs (miRNAs) in the hippocampus. Notably, these alterations were associated with cognitive impairments induced by binge-like ethanol treatment in adolescent mice (Montesinos et al., 2015; Pascual et al., 2021; Mellado et al., 2023).

## MATERIALS AND METHODS

### MSC isolation, culture and isolation of MSC-EVs

Human adipose tissue was obtained from surplus fat tissue during knee prosthesis operation performed on four patients under sterile conditions. The human samples were anonymized. The experimental procedure had been previously evaluated and accepted by the Regional Ethics Committee for Clinical Research with Medicines and Health Products following Code of Practice 2014/01. As exclusion criteria, no samples were collected from patients with a history of cancer or infectious diseases at the time of (viral or bacterial) surgery. All the human patients voluntarily signed an informed consent document to allow use of the adipose samples. Cells were expanded and grown in growth medium (GM: High glucose DMEM basal medium supplemented with 20% FBS [previously centrifuged at 100000 x g for 1 h for EV depletion and then filtered by a 0.2 μm filter], 100 units/ml penicillin, 100 µg/ml streptomycin and 2 mM L-glutamine). MSCs have been characterized and previously described (Mellado-López et al., 2017; Muñoz-Criado et al., 2017). Subconfluent cells were incubated in GM for 48 h. Then media were collected and cleared from detached cells and cell fragments by centrifugation at 300 x g for 10 min, and by the supernatant at 2000 x g for 10 min, respectively. Subsequently, apoptotic bodies and other cellular debris were pelleted by centrifuging the resulting supernatant at 10000 x g for 30 min. EVs were then pelleted from the previous resulting supernatant at 100000 x g for 1 h. The EVs pellet was washed with PBS and centrifuged at 100000 x g for 1 h. EVs were finally suspended in 100 ml PBS and stored at −80 °C.

### Animals and treatments

Thirty-two female C57BL/6 WT mice (Harlan Ibérica, Barcelona, Spain) were used. Mice were housed (3-4 animals/cage) and maintained on a water and solid diet *ad libitum*. Environmental conditions, such as light and dark cycles (12/12 h), temperature (23 °C), and humidity (60%), were controlled for all the animals. All the experimental procedures were carried out in accordance with the guidelines approved by the European Communities Council Directive (86/609/ECC) and Spanish Royal Decree 53/2013 with the approval of the Ethical Committee of Animal Experimentation of the Príncipe Felipe Research Centre (Valencia, Spain) and the Generalitat Valenciana on August 8th, 2020 (Project identification code: 2020/VSC/PEA/0145).

The intermittent ethanol treatment was initiated early in adolescence or during the prepubescent period on postnatal day (PND) 30 (Brust et al., 2015). Morning doses of either saline or 25% (v/v) ethanol (3 g/kg) in isotonic saline were administered intraperitoneally to 30-day-old mice on 2 consecutive days with 2-day gaps without injections for 2 weeks (PND30 to PND43), as previously described (Pascual et al., 2007; Montesinos et al., 2015). No signs of peritoneal cavity irritation, pain or distress or peripheral inflammation induced by intraperitoneal ethanol concentration were noted, which agrees with other studies that have used intraperitoneal ethanol administration (Allen-Worthington et al., 2015). After a single ethanol dose, blood alcohol levels peaked at 30 min (∼340 mg⁄dL) and then progressively lowered for 5 hours post-injection. Three hours before ethanol administration, animals were also treated with MSC-EVs (50 µg/dose) or saline (sodium chloride, 0.9%) in the tail vein once a week (with the third and seventh ethanol dose). Animals were randomly assigned to four groups according to their treatments: 1) physiological saline or control, 2) physiological saline plus MSC-EVs, 3) ethanol, and 4) ethanol plus MSC-EVs. No changes in either animals’ body weight or brain weight were observed during the intermittent treatment (Mellado et al., 2023). Animals were anesthetized 24 hours after the last (8th) ethanol or saline administration (PND 44). Brains were removed and transferred to a plate placed on ice. Olfactory bulbs, the cerebellum and the pons were removed. Brains were placed with the ventral side facing the plate and both hemispheres were separated. Then, we performed the hippocampus dissection with two spatulas, separating the cortex overlying the dorsal hippocampus (Sultan, 2013; Harris et al., 2019; Le Merre et al., 2021; Jaszczyk et al., 2022). The hippocampi (n=8 mice/group) were immediately snap-frozen in liquid nitrogen and stored at −80 °C until used.

### MSC-EVs characterization by transmission electron microscopy and nanoparticles tracking analysis

The freshly isolated EVs were fixed with 2% paraformaldehyde and prepared as previously described (Ibáñez et al., 2019). Preparations were examined under a transmission FEI Tecnai G2 Spirit electron microscope (FEI Europe, Eindhoven, the Netherlands) with a Morada digital camera (Olympus Soft Image Solutions GmbH, Münster, Germany). In addition, an analysis of the absolute size range and concentration of microvesicles was performed using NanoSight NS300 Malvern (NanoSight Ltd., Minton Park, UK), as previously described (Ibáñez et al., 2019). Figure 1 (A, C) reports the characterization of EVs by electron microscopy and nanoparticle tracking analysis.

**Figure 1.**
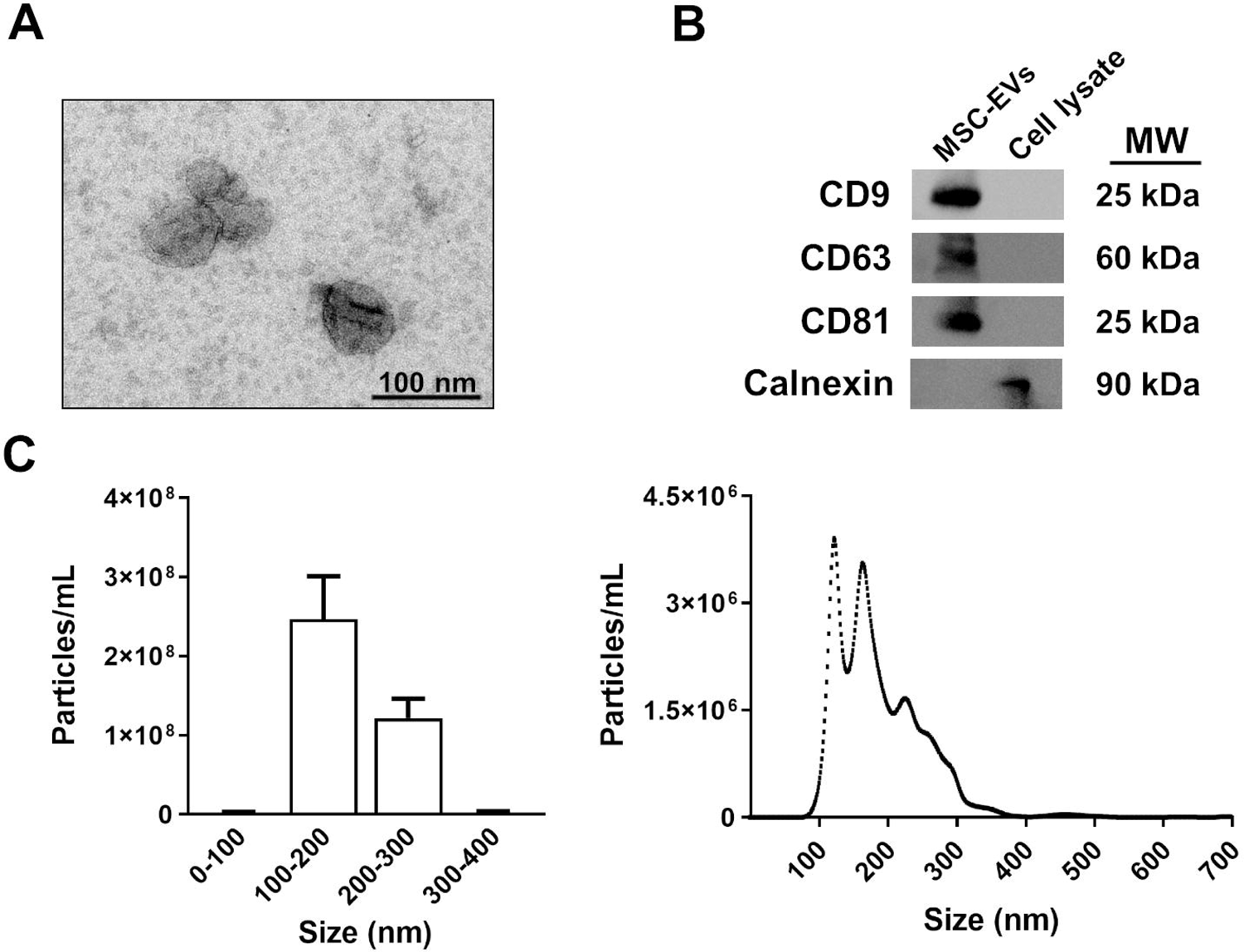
Characterization of MSC-EVs. **(A)** Electron microscopy image of MSC-EVs. **(B)** Analysis of the protein expression of EV markers (CD9, CD63, and CD81) in EVs and cell lysates. Calnexin expression was used to discount cytosolic protein contamination in EV samples. Cell lysates from astroglial cells were used as a positive control for calnexin expression. A representative immunoblot for each protein is shown. **(C)** Measurement of the size distribution and concentration of MSC-EVs by the nanoparticles tracking analysis. A high peak ranging between 100 and 200 nm is shown, which includes the size range of EVs.

### Western blot analysis

The Western blot technique was performed in MSC-EVs for their characterization purposes (Fig. 1B), as previously described (Montesinos et al., 2015). The employed primary antibodies were: anti-CD9, anti-CD63, anti-CD81 and anti-calnexin (Santa Cruz Biotechnology, USA). Membranes were washed, incubated with the corresponding HRP-conjugated secondary antibodies and developed by the ECL system (ECL Plus; Thermo Scientific, Illinois, USA). The cell lysate from the astrocyte primary cell culture was used as the negative control for CD9, CD63 and CD81, and as the positive control for calnexin. The full unedited blots are included in the Supplementary Material (Fig. S1).

### RNA isolation, reverse transcription and quantitative RT-PCR

The frozen hippocampal samples were used for total RNA extraction. Tissues were disrupted using TRIzol (Sigma-Aldrich, St. Louis, USA), and the total RNA fraction was extracted following the manufacturer’s instructions. Total mRNA and total miRNA were reverse-transcribed by the NZY First-Strand cDNA Synthesis Kit (NZYTech, Lda. Genes and Enzymes, Lisboa, Portugal) and TaqMan^TM^ Advanced miRNA Assays (ThermoFisher Scientific, USA), respectively.

RT-qPCR was performed in a QuantStudio^TM^ 5 Real-Time PCR System (Applied Biosystems, Massachusetts, USA). Genes were amplified employing the AceQ® qPCR SYBR Green Master Mix (NeoBiotech, Nanterre, France) following the manufacturer’s instructions. The mRNA level of housekeeping gene cyclophilin A was used as an internal control for the normalization of the analyzed genes. Specific miRNAs assays were amplified by the TaqMan^TM^ Fast Advanced Master Mix (ThermoFisher Scientific) and snRNA U6 was used as an internal control. All the RT-qPCR runs included non-template controls (NTCs). Experiments were performed in triplicates. Quantification of expression (fold change) from the Cq data was calculated by the ΔΔCq method (Schmittgen and Livak, 2008) by the QuanStudio^TM^ Design & Analysis Software (Applied Biosystems). Details of the nucleotide sequences of the used primers and miRNAs assays are detailed in the Supplementary Material (Tables S1 and S2).

### Bioinformatic analysis of miRNAs

Based on the three miRNAs of interest (mmu-miR-21a-5p, mmu-miR-141-5p, mmu-miR-146a-5p), we used the multiMiR package and database (version 1.16.0) (Ru et al., 2014) to obtain their validated target genes. Subsequently, we conducted an overrepresentation analysis using the ClusterProfiler package (version 4.2.2) (Wu et al., 2021) to identify the KEGG pathways (Kanehisa and Goto, 2000) that are enriched in the list of target genes, compared to what would be expected by chance. We then compared the results for each miRNA to evaluate the similarities. The R version used for this analysis was 4.1.2 [R Core Team (2021). R: A language and environment for statistical computing. R Foundation for Statistical Computing, Vienna, Austria. https://www.R-project.org/]. Protein-protein interaction (PPI) networks were also constructed using the STRING web tool for the target-shared genes among the three miRNAs (Szklarczyk et al., 2023). The PPI enrichment was assessed using a minimum required interaction score of “high confidence” (0.7).

### Statistical analysis

The results are reported as the mean±SEM. All the statistical parameters used were calculated with SPSS v28. The Shapiro-Wilk test was used to test for data distribution normality. A one-way ANOVA was used as parametric tests, followed by the Bonferroni post hoc test. The Kruskal-Wallis test was used as a non-parametric alternative. Values of p < 0.05 were considered statistically significant.

## RESULTS

### MSC-EVs restore the activation of the NLRP3 inflammasome pathway in the hippocampus of adolescent mice treated with binge-like ethanol treatment

Our previous results demonstrated that MSC-EVs from adipose tissue were able to ameliorate the neuroinflammation induced by binge drinking in adolescent mice (Mellado et al., 2023). Considering that ethanol can activate the NLRP3 inflammasome in astroglial cells and the cerebral cortices of mice (Alfonso-Loeches et al., 2014, 2016), we therefore wondered if an intravenous injection of MSC-EVs was also capable of decreasing the activation of the NLRP3 inflammasome pathway in the hippocampus of ethanol-treated adolescent mice. To answer this question, we first evaluated the components of the NLRP3 inflammasome complex, which requires the adaptor protein ASC and the caspase-1 to activate cytokine release (Almeida-da-Silva et al., 2023). Figure 2A shows that the ethanol treatment increases the gene expression of NLRP3 [F(3,28) = 2.103, p < 0.0001], caspase-1 [F(3,27) = 0.5897, p < 0.0001] and ASC [F(3,27) = 0.8906, p = 0.0147], compared to the saline-treated adolescents. Analyzing the cytokine expression, ethanol was also capable of increasing the mRNA levels of IL-1β [F(3,27) = 4.358, p = 0.0056] and IL-18 [F(3,28) = 2.534, p = 0.0024], compared to the saline group (Fig. 2B). Notably, the administration of MSC-EVs was able to attenuate the ethanol-induced mRNA expression of the NLRP3 inflammasome complex [NLRP3 [F(3,28) = 2.103, p = 0.0001], caspase-1 [F(3,27) = 0.5897, p = 0.0005] and ACS [F(3,27) = 0.8906, p = 0.0025]] (Fig. 2A) and the cytokine levels of IL-1β [F(3,27) = 4.358, p < 0.0001] and IL-18 [F(3,28) = 2.534, p = 0.0023] (Fig. 2B). These data also showed statistical significant differences between animals treated with saline+MSC-EVs and ethanol in the gene expression of NLRP3 [F(3,28) = 2.103, p < 0.0001], caspase-1 [F(3,27) = 0.5897, p = 0.0001], ACS [F(3,27) = 0.8906, p < 0.0207], IL-1β [F(3,27) = 4.358, p = 0.0014] and IL-18 [F(3,28) = 2.534, p < 0.0001].

**Figure 2.**
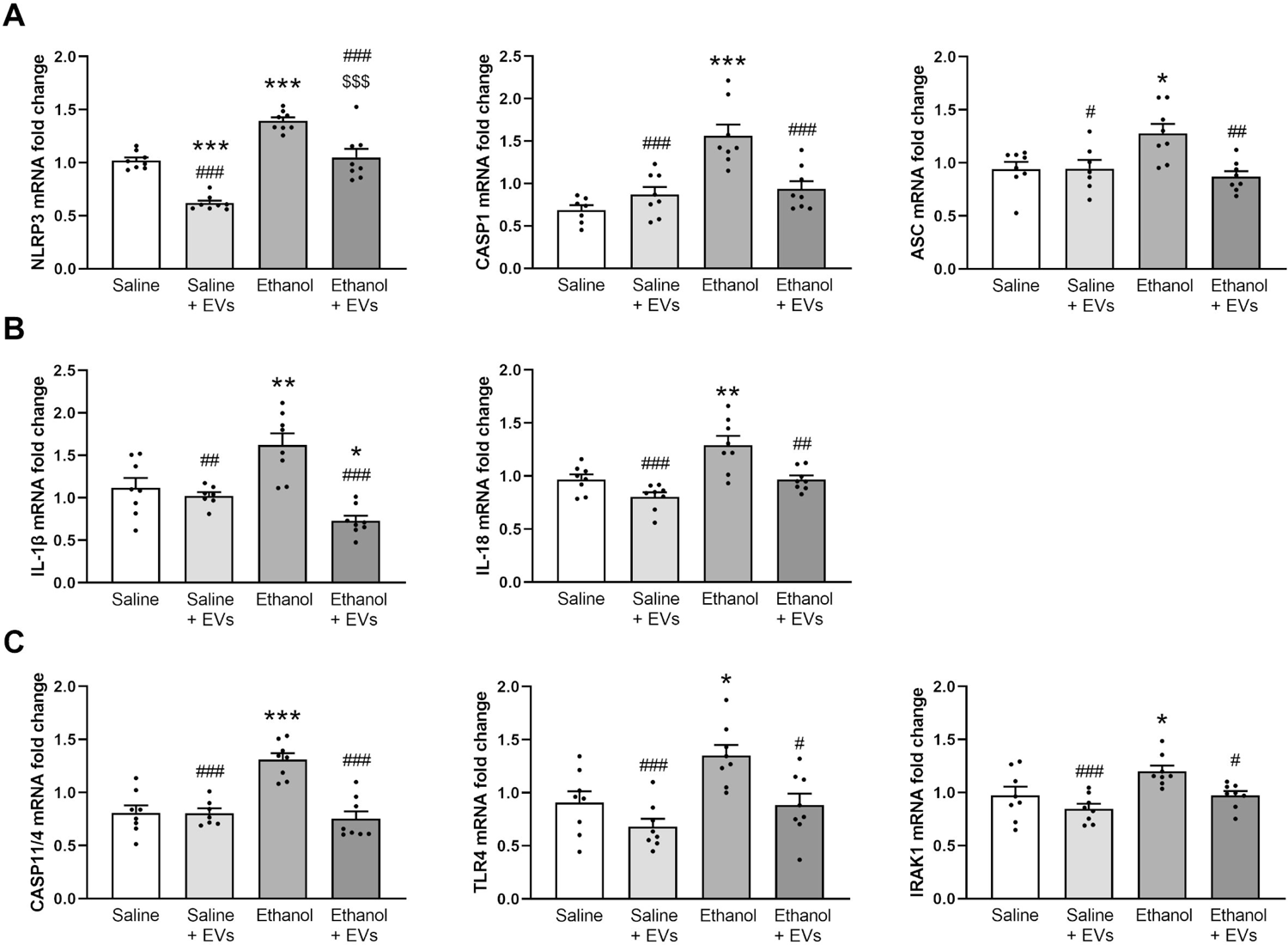
MSC-EVs attenuate the activation of hippocampal NLRP3 inflammasome complex induced by binge-like ethanol treatment in adolescent mice. **(A)** The mRNA levels of components of the NLRP3 inflammasome complex, such as NLRP3, caspase-1 (CASP-1) and ASC in the hippocampus. **(B)** The mRNA expression of the proinflammatory IL-1β and IL-18. **(C)** The mRNA levels of the genes, caspase-11/4 (CASP11/4), TLR4 and IRAK1. Data represent mean±SEM, n=8 mice/group. * p < 0.05, ** p < 0.01 and *** p < 0.001, compared to their respective saline-treated group; #p < 0.05, ## p < 0.01 and ### p < 0.001, compared to their respective ethanol-treated group; $$$ p < 0.001 compared to their respective saline+MSC-EVs treated group.

Considering that several inflammatory molecules, such as caspase-11/4, TLR4 or IRAK1 may activate the NLRP3/caspase-1 complex (Kelley et al., 2019; Zheng et al., 2020), we then evaluated the gene expression of these genes (Fig. 2C). The results shown, that while ethanol treatment increased the levels of caspase-11/4 [F(3,27) = 0.3200, p < 0.0001], TLR4 [F(3,28) = 0.3609, p = 0.0181] and IRAK1 [F(3,28) = 1.592, p = 0.0438], the administration of MSC-EVs was able to restore the upregulation I of these genes (caspase-11/4 [F(3,27) = 0.3200, p < 0.0001], TLR4 [F(3,28) = 0.3609, p = 0.0122] and IRAK1 [F(3,28) = 1.592, p = 0.0446]) in ethanol-treated mice. Significant differences were also shown between the animals treated with saline+MSC-EVs and ethanol [caspase-11/4 [F(3,27) = 0.3200, p < 0.0001], TLR4 [F(3,28) = 0.3609, p = 0.0003] and IRAK1 [F(3,28) = 1.592, p < 0.0009]].

To gain further insight into how ethanol affects inflammasome activation and which type of NLRs was activated in the hippocampus of ethanol-treated adolescent mice, the gene expression of other NLRs (e.g., NLRP1, NLRC4, and AIM2) were also assessed. Figure 3 shows that whereas ethanol treatment upregulated the expression of NLRP1 [F(3,28) = 0.6065, p = 0.0023], AIM2 [F(3,28) = 0.4111, p = 0.0119] and NLRC4 [F(3,28) = 3.775, p = 0.0027], the intravenous injection of MSC-EVs was capable of attenuating these increases in ethanol-treated adolescent mice [NLRP1 [F(3,28) = 0.6065, p = 0.0209], AIM2 [F(3,28) = 0.4111, p = 0.0472] and NLRC4 [F(3,28) = 3.775, p = 0.0407]]. We noted significant differences between the animals treated with saline+MSC-EVs and ethanol [NLRP1 [F(3,28) = 0.6065, p = 0.0021], AIM2 [F(3,28) = 0.4111, p = 0.0072] and NLRC4 [F(3,28) = 3.775, p = 0.0024]].

**Figure 3.**
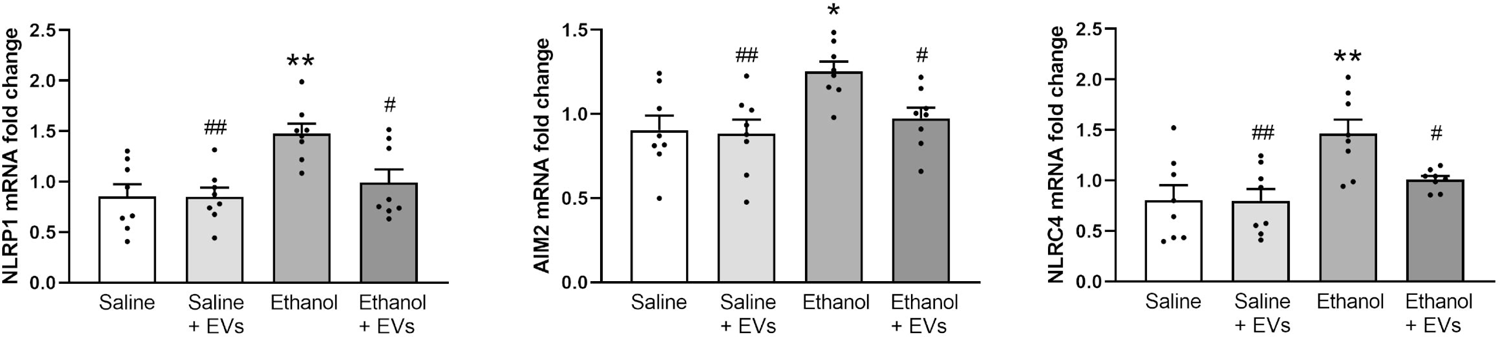
MSC-EVs diminish the activation of other NLRs induced by binge-like ethanol treatment in adolescent mice. The mRNA expression of the NLRs, NLRP1, AIM2 and NLRC4 in the hippocampus. Data represent mean±SEM, n=8 mice/group. * p < 0.05, ** p < 0.01 and *** p < 0.001, compared to their respective saline-treated group; # p < 0.05, ## p < 0.01 and ### p < 0.001, compared to their respective ethanol-treated group.

### MSC-EVs ameliorate hippocampal inflammatory genes and miRNAs induced by binge-like ethanol treatment in adolescent mice

We then analyzed if MSC-EVs administration could restore the up-regulation in the mRNA levels of inflammatory genes induced by ethanol treatment. We measured the iNOS, NF-κB, MCP-1 and CX3CL1 levels in the hippocampus under different experimental conditions. Figure 4 shows that ethanol treatment increased the gene expression of iNOS [F(3,28) = 1.481, p = 0.0051], NF-κB [F(3,28) = 0.09025, p = 0.0116], MCP-1 [F(3,28) = 3.211, p = 0.0145], and CX3CL1 [F(3,28) = 1.159, p = 0.0019], when compared to their saline counterparts. As expected, MSC-EVs administration was able to attenuate the ethanol-induced expression of these proinflammatory molecules compared to the ethanol-treated mice [iNOS [F(3,28) = 1.481, p = 0.0411], NF-κB [F(3,28) = 0.09025, p = 0.0195], MCP-1 [F(3,28) = 3.211, p = 0.0010], and CX3CL1 [F(3,28) = 1.159, p = 0.0002]]. In addition, ethanol treatment significantly increased the iNOS [F(3,28) = 1.481, p = 0.0013], NF-κB [F(3,28) = 0.09025, p = 0.0006], MCP-1 [F(3,28) = 3.211, p = 0.0007], and CX3CL1 [F(3,28) = 1.159, p < 0.0001] levels compared to the saline plus MSC-EVs-treated mice. No changes in the expression of these genes were found between the saline- and MSC-EVs-treated animals.

**Figure 4.**
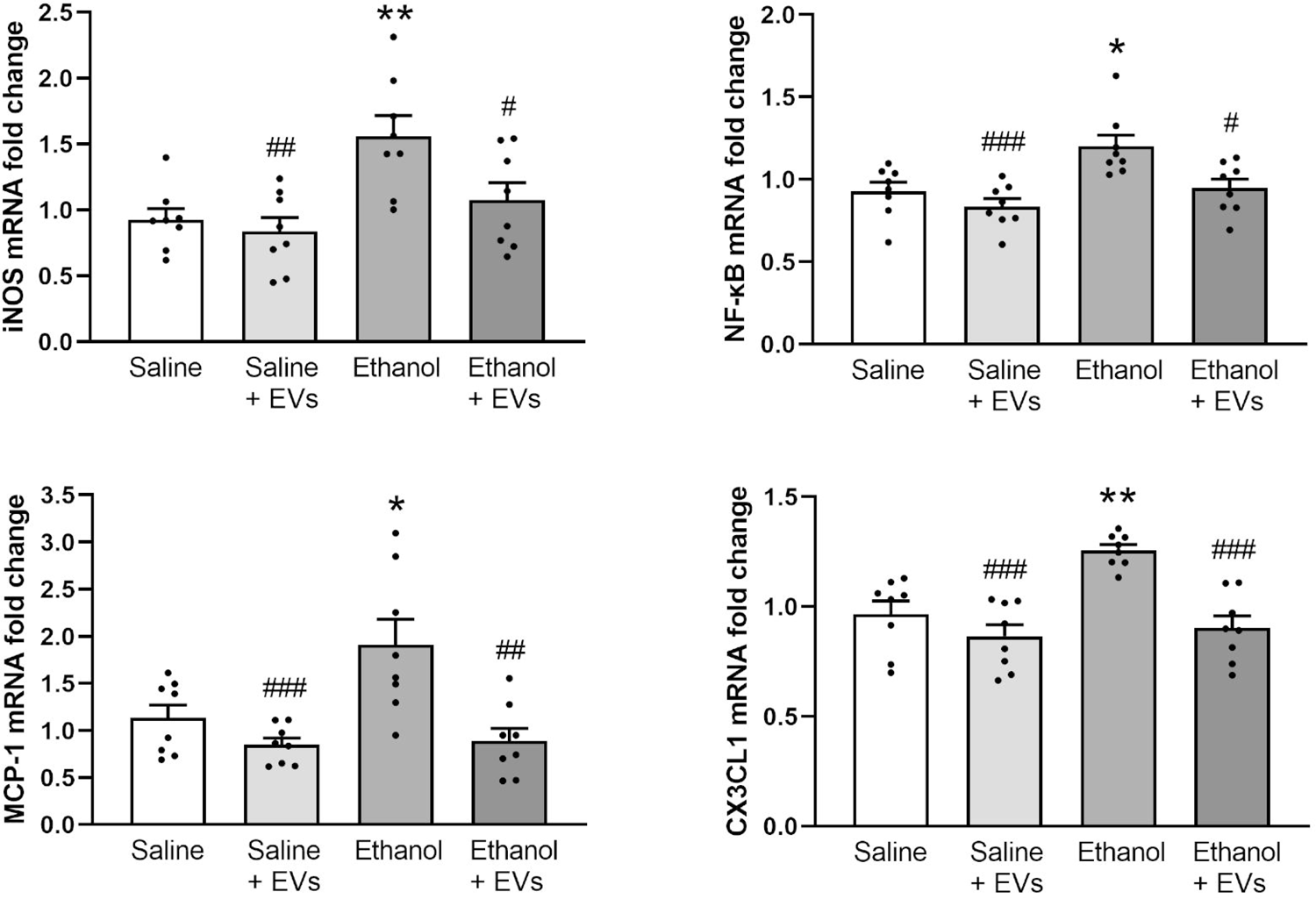
MSC-EVs decrease the levels of inflammatory genes induced by binge-like ethanol treatment in adolescent mice. The mRNA levels of iNOS, NF-κB, MCP-1, and CX3CL1 were analyzed in the hippocampal samples. Data represent mean±SEM, n=8 mice/group. * p < 0.05 and ** p < 0.01, compared to their respective saline-treated group; # p < 0.05, ## p < 0.01 and ### p < 0.001, compared to their respective ethanol-treated group.

We then wondered whether the alterations observed in the expression of the inflammatory genes (Fig. 2-4) in the hippocampus of ethanol-treated adolescent mice could be associated with the changes in the expression of several miRNAs, such as miR-21a-5p, miR-141-5p and miR-146a-5p (Fig. 5). Indeed, our studies demonstrate the action of ethanol on the miRNA profiles (Ureña-Peralta et al., 2018; Ibáñez et al., 2019). Notably, a significant increase in the miR-21a-5p [F(3,27) = 3.749, p = 0.0001], miR-141-5p [F(3,27) = 1.424, p = 0.0398] and miR-146a-5p [F(3,26) = 3.522, p < 0.0026] levels took place in the adolescent mice treated with ethanol. This expression significantly decreased in the animals treated with ethanol plus MSC-EVs compared to the ethanol-treated mice [miR-21a-5p [F(3,27) = 3.749, p < 0.0001], miR-141-5p [F(3,27) = 1.424, p = 0.0008] and miR-146a-5p [F(3,26) = 3.522, p = 0.0002]]. Saline plus MSC-EVs-treated mice showed lower levels of expression compared with the ethanol-treated group [miR-21a-5p [F(3,27) = 3.749, p < 0.0001], miR-141-5p [F(3,27) = 1.424, p = 0.0006] and miR-146a-5p [F(3,26) = 3.522, p = 0.0002]].

**Figure 5.**
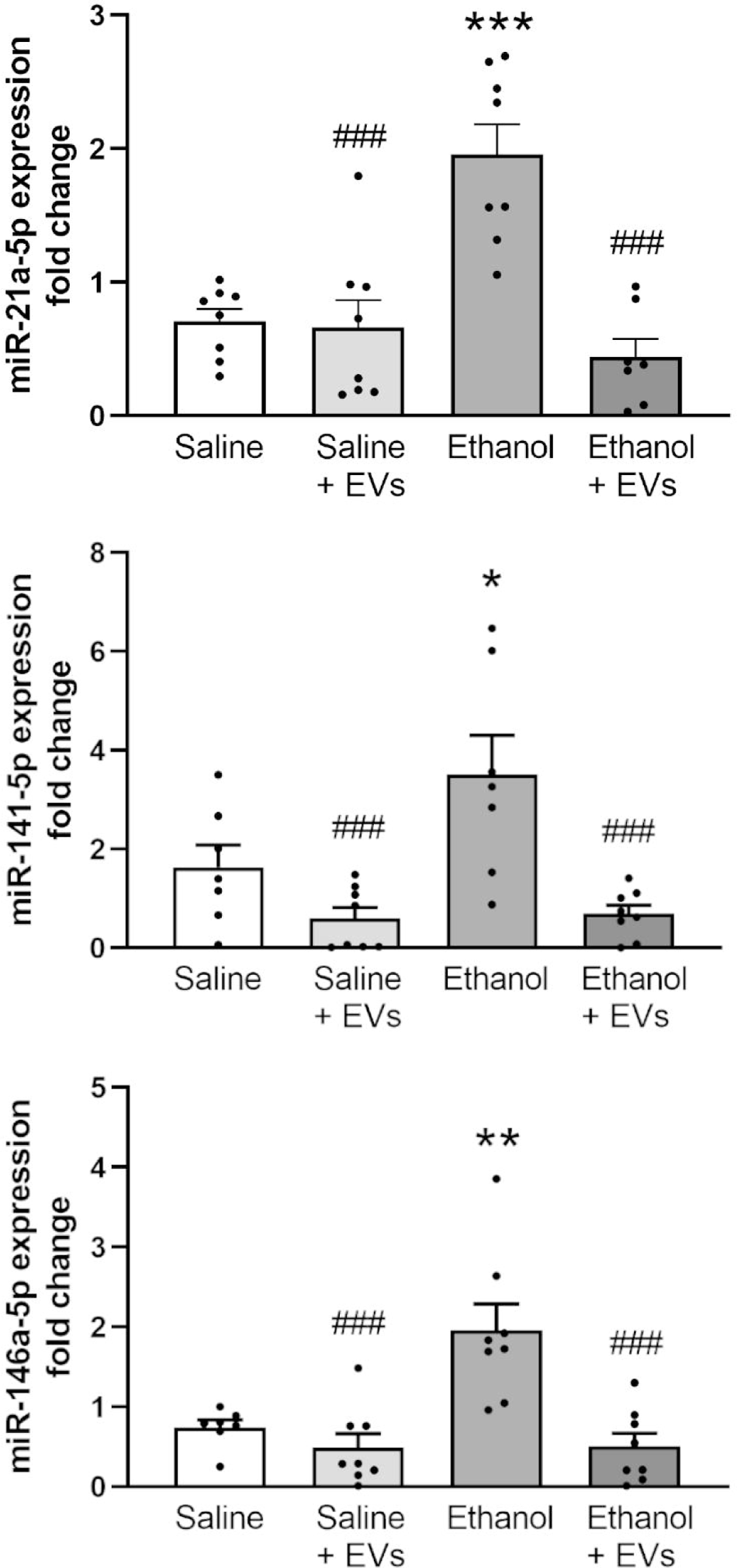
MSC-EVs restore the levels of miR-21a-5p, miR-141-5p and miR-146a-5p in the hippocampus of binge-like ethanol-treated adolescent mice. Data represent mean±SEM, n=8 mice/group. * p < 0.05, ** p < 0.01 and *** p < 0.001, compared to their respective saline-treated group; ### p < 0.001, compared to their respective ethanol-treated group.

### Bioinformatic analysis of miRNAs involved in the effects of ethanol and MSC-EVs

After confirming that MSC-EVs are able to restore the ethanol-induced alterations in the levels of some miRNAs (miR-21a-5p, miR-146a-5p and miR-141-5p), we performed a functional analysis for these noncoding RNAs to further establish the main biological functions and key pathways involved in the effects of ethanol and MSC-EVs. The bioinformatic analysis was separated into different steps. The first step determined the preferential targets modulated by each miRNA using the multiMiR package and database (version 1.16.0) (Ru et al., 2014). The analysis demonstrated that the 2205, 397 and 4 target genes were modulated by miR-21a-5p, miR-146a-5p and miR-141-5p, respectively (data not shown). Figure 6A reveals that miR-21a-5p and miR-146a-5p displayed a stronger interaction with the modulated target genes than the other two intersections, which had 0 or 1 shared genes. For instance, miR-21a-5p and mir-146a-5p control immune target genes, such as Irak2, Ccr5, Cxcr4, Irf4, Tirap, Cd28 and Camk2d, as well as a gene required for RNA-mediated gene expression named Ago2. Using the STRING web tool, we created PPI networks for the 119 miRNA-targeted shared genes (Fig. 6A). Within the PPI analysis, we obtained three larger clusters and six with almost one connection (Fig. 6B). The network had significantly more interactions than expected (PPI enrichment p-value: 9.09e-06), indicating that this group of proteins has more interactions among themselves than what would be expected for a random set of proteins of the same size and degree distribution drawn from the genome. Such enrichment suggests that these proteins are at least partially biologically connected as a group. Afterwards, we conducted an overrepresentation analysis using the KEGG pathways, which allowed us to identify the signaling pathways enriched in the list of all the target genes. Figure 6C shows the significantly enriched KEGG pathways obtained by miRNA and the number of miRNA-regulated genes that participate in the significantly enriched KEGG pathways for each miRNA. Although the three miRNAs are able to modulate signaling pathways associated with the immune response and inflammatory processes (e.g., TLR, cytokine receptor, chemokine, B cell receptor TGF-β, MAPK, etc.), the miR-21a-5p is able to modify the NOD-like receptor signaling pathway.

**Figure 6.**
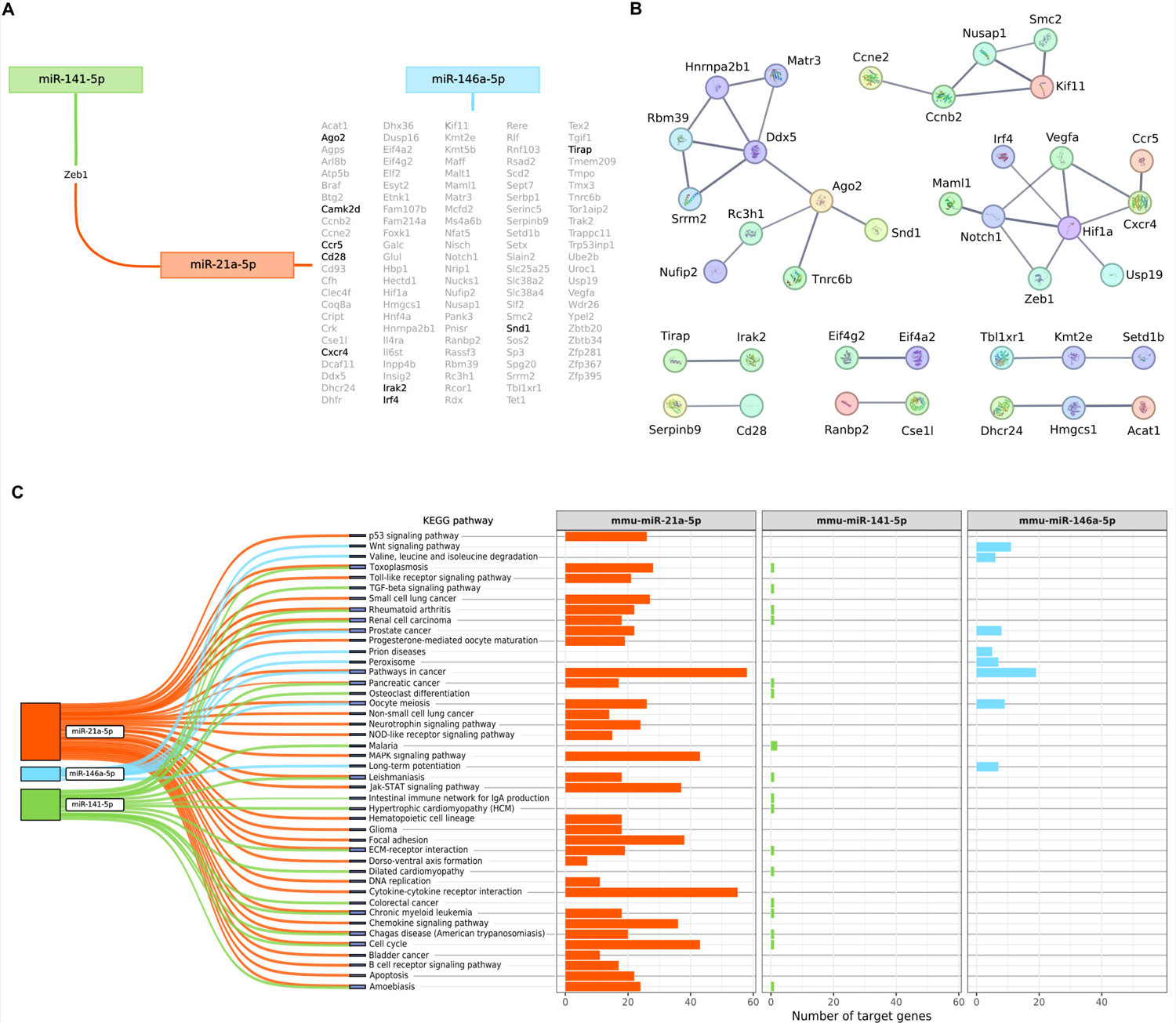
Main results of bioinformatics analysis focused on the miRNAs miR-21a-5p, miR-141-5p, miR-146a-5p. **(A)** Target genes shared by the miRNAs. Genes highlighted in black are related to processes involved in the immune response and RNA-mediated gene expression. **(B)** Protein-protein interaction (PPI) networks generated from all the miRNA-targeted shared genes. Line thickness indicates the strength of data support. **(C)** Number of miRNA-targeted genes that participate in the significantly enriched KEGG pathways for each miRNA.

## DISCUSSION

We have recently demonstrated the therapeutic role of MSC-EVs to restore neuroinflammation, myelin and synaptic structural alterations in the prefrontal cortex, as well as cognitive dysfunctions induced by binge-like ethanol treatment in adolescent mice (Mellado et al., 2023). Considering the involvement of the inflammasome receptors in the neuroinflammatory processes (Piancone et al., 2021) and behavioral dysfunctions (Jin et al., 2019; Hou et al., 2020), the present findings provide evidence that the intravenous administration of MSC-EVs restores the activation of the NLRP3 inflammasome complex and other NLRs inflammasomes (e.g., NLRP1, NLRC4 and AIM2), as well as the alterations of inflammatory genes and miRNAs in the hippocampus induced by binge-like ethanol treatment in adolescent mice (Fig. 7).

**Figure 7.**
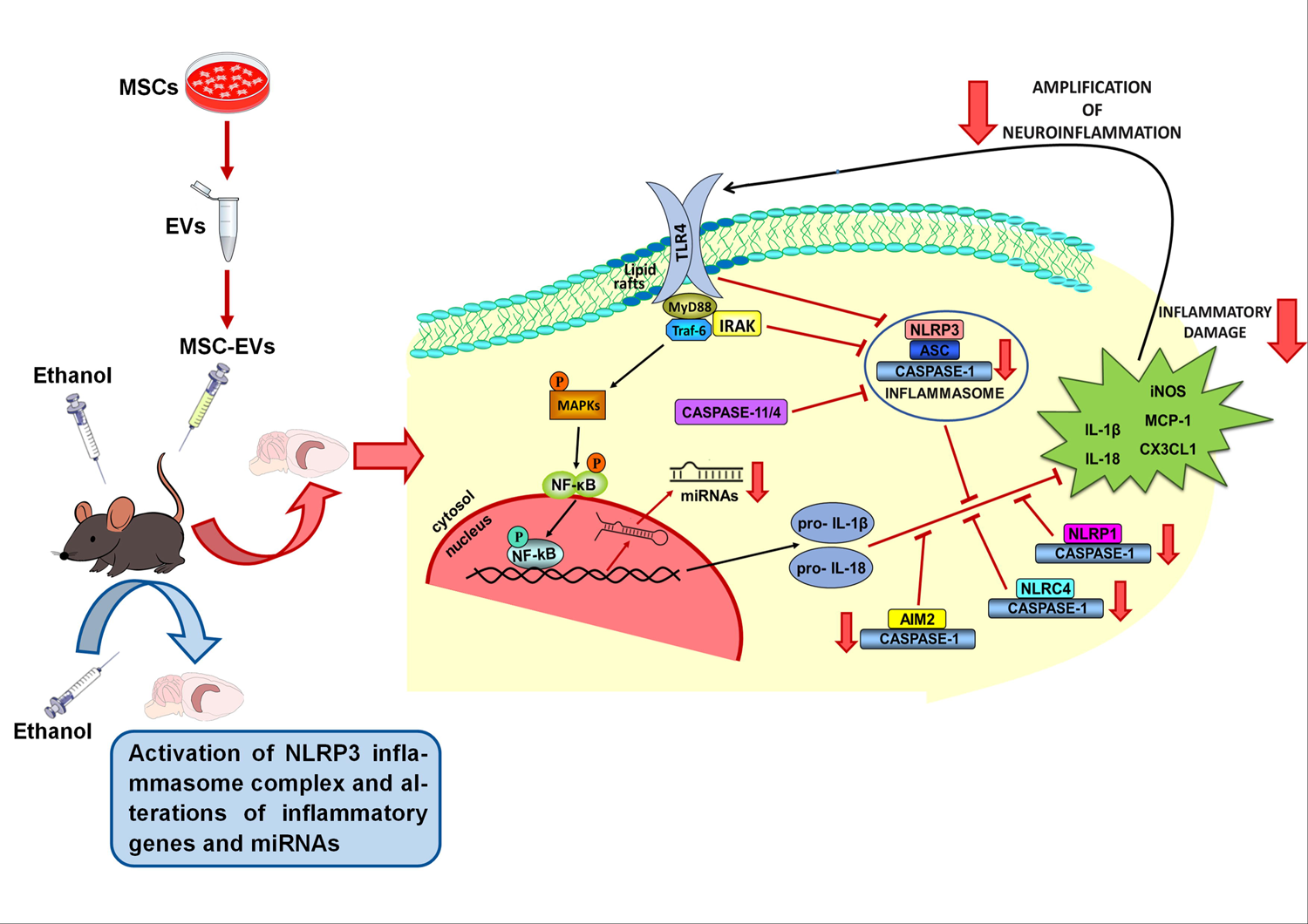
Schematic representation of the protective effects of MSC-EVs administration on ethanol-induced hippocampal neuroinflammation, by inhibiting NLRP3 inflammasome activation. MSC-EVs from adipose tissue were administered by intravenous injection (iv) prior to ethanol treatment (ip, intraperitoneal injection) in adolescent mice. MSC-EVs administration ameliorates the activation of the hippocampal NLRP3 inflammasome complex and other NLRs inflammasomes (e.g., NLRP1, NLRC4 and AIM2), as well as the alterations in the expression of inflammatory genes and miRNAs induced by binge-like ethanol treatment in adolescent mice.

Our previous studies revealed the role of NLRP3/caspase-1 complex activation, along with the up-regulated pro-inflammatory cytokines and chemokines levels in ethanol-treated astrocytes and in cortices from chronic alcohol-fed WT mice (Alfonso-Loeches et al., 2014, 2016). Likewise, we have also provided evidence that binge-ethanol treatment in adolescent mice activates NLRs and their signaling pathways, inducing a neuroinflammatory response. Since the MSC-EVs from adipose tissue were capable of diminishing the prefrontal cortex neuroinflammation induced by binge drinking in adolescent mice (Mellado et al., 2023), the use of these microvesicles could also participate in ameliorating the inflammasome complex in the hippocampus. In this line, it has been reported that the administration of MSC-EVs alleviates the inflammatory response by inhibiting the activation of NLRP3 inflammasome in an Alzheimer’s disease mouse model (Xu et al., 2022), in spinal cord injury (Noori et al., 2021) or in lipopolysaccharide (LPS)-induced cardiomyocyte inflammation (Pan et al., 2022). The present study provide evidence that MSC-EVs restores the activation, not only of the hippocampal NLRP3 inflammasome complex, but also of up-regulated inflammatory genes (IL-1β, IL-18, iNOS, NF-κB, MCP-1 and CX3CL1) and miRNAs (miR-21a-5p, miR-141-5p and miR-146a-5p) in the hippocampus of adolescent mice with binge-like ethanol treatment. Indeed, the bioinformatic analysis of miR-21a-5p and miR-146a-5p revealed the involvement of these miRNAs with inflammatory target genes and NOD-like receptor signaling pathways.

In the last few years, a large number of studies reported the therapeutical effects of miRNAs contained within MSC-EVs. These therapeutical effects have been attributed to their actions in regulating the expression of multiple target genes and in modulating various cell signaling processes to promote functional recovery (Qiu et al., 2018). Dysregulation of circulating levels of specific EV-miRNAs has been reported in patients with neurodegenerative diseases. Several EV-miRNAs (e.g., miR-124a, miR-146a, miR-21, miR-29b, miR-873a-5p) have been reported to show potential effects to reduce oxidative stress and neuroinflammation in Alzheimer’s disease or traumatic brain injury models (Bang and Kim, 2022). Another study has also demonstrated the protective function of these microvesicles by transferring the miR-223-3p to suppress the circular RNA PWWP2A, thereby alleviating pulmonary fibrosis through the NLRP3 signaling pathway (Hou et al., 2023). Conversely, our results show that MSC-EVs are able to diminish the levels of the inflammatory miRNAs (miR-21a-5p, miR-146a-5p and miR-141-5p) induced by binge-like ethanol treatment in adolescent mice.

Accordingly, a TLRs inflammatory response, associated with the upregulation of the miR-132, can be decreased through this miRNA located within MSC-EVs by increasing the IL-10 expression and decreasing NF-κB and IL-1β levels (Hade et al., 2021). Although miRNAs can modulate gene expression, the transcription of these miRNAs can be regulated by additional molecules (e.g., Argonaute protein, Drosha or DGCR8), which can enhance or inhibit this process, modifying the cell’s fate (Ergin and Çetinkaya, 2022). Therefore, we suggest that EV-mediated transfer of miRNAs and other molecules between MSCs and neural cells, may be involved in promoting functional recovery in ethanol-treated adolescent mice.

The hippocampus is an important brain area associated with memory and learning processes (Gandhi et al., 2014; Pascual et al., 2021). Therefore, the involvement of the MSC-EVs to restore the alterations in the NLPR3 inflammasome induced by binge-like ethanol treatment in the adolescent hippocampus could also participate in the ethanol-induced cognitive dysfunction shown in Mellado et al. (2023). Several studies have reported that spatial memory dysfunction could be related to the neuroinflammation induced by the hippocampal activation of the NLRP3 inflammasome and the TLR4/NF[κB signaling pathway in a mouse model of Alzheimer’s disease (Jin et al., 2019) and in an experimental autoimmune encephalomyelitis (Hou et al., 2020).

Our interest in this study was to provide the therapeutic role of MSC-EVs from adipose tissue to restore the NLPR3 neuroinflammatory response induced by binge drinking in adolescence. Although adipose tissue is one of the best alternatives as a source for MSCs due to the accessibility and abundance of this tissue, some limitations can also be presented in this study. For instance, 1) we designed the term EVs, since we cannot distinguish between exosomes and microvesicles, which contain similar size and protein markers, 2) the miRNA profile of MSC-EV differs depending on the origin tissue, and 3) we cannot discard the possibility of other miRNAs or molecules involved in the neuroprotective effects.

Taken together, these results support for the first time the protective role of MSC-EVs in the hippocampal neuroinflammatory immune response through the activation of the NLRP3 inflammasome complex induced by binge-like ethanol treatment in adolescence. This study also evidences a possible mechanism of action in the effects of ethanol and MSC-EVs in the inflammasome alterations through the miR-21a-5p and mir-146a-5p regulation. In conclusion, the present findings suggest the therapeutic action of MSC-EVs as a new tool to restore the neuroinflammatory response related with alcohol consumption and brain damage.

## ACKNOWLEDGMENTS

This work has been supported by grants from the Spanish Ministry of Health[PNSD (2019[I039), by FEDER/Ministerio de Ciencia e Innovación – Agencia Estatal de Investigación PID2021-1243590B-I100, GVA (CIAICO/2021/203), the Primary Addiction Care Research Network (RD21/0009/0005) and FEDER Funds, GVA.

## CONFLICT OF INTEREST

All authors declare no competing interest in research conduction and paper writing.

## AUTHOR CONTRIBUTIONS

MP, SM and CG conceived and designed the experiments. SM, MJMB and CPC performed the experiments and analyzed the data. SM, MJMB, CPC, CG and MP wrote the manuscript. SM, FGG, VMM, CG and MP revised the manuscript. All authors read and approved the final manuscript.

## DATA AVAILABILITY STATEMENT

The data that support the findings of this study are available on request from the corresponding author. The data are not publicly available due to privacy or ethical restrictions.

## ABBREVIATIONS

CAMK2D: calcium/calmodulin dependent protein kinase II delta

CCR5: C-C chemokine receptor type 5

CD: Cluster of Differentiation

CX3CL1: C-X3-C motif chemokine ligand 1 or fractalkine

CXCR4: C-X-C motif chemokine receptor 4

DGCR8: DiGeorge Critical Region 8

iNOS: inducible nitric oxide synthase

IRAK: interleukin-1 receptor-associated kinase

IRF4: interferon regulatory factor 4

MAPK: mitogen-activated protein kinase

MCP-1: monocyte chemoattractant protein-1

NF-κB: nuclear factor-kappa B

TGF-β: Transforming growth factor beta

TIRAP: Toll-interleukin-1 Receptor (TIR) domain-containing adaptor protein.

## Notes

### Competing Interest Statement

The authors have declared no competing interest.

## REFERENCES

Alfonso-Loeches S, Pascual-Lucas M, Blanco AM, Sanchez-Vera I, Guerri C. 2010. Pivotal role of TLR4 receptors in alcohol-induced neuroinflammation and brain damage. J Neurosci Off J Soc Neurosci 30:8285–8295.

Alfonso-Loeches S, Ureña-Peralta J, Morillo-Bargues MJ, Gómez-Pinedo U, Guerri C. 2016. Ethanol-Induced TLR4/NLRP3 Neuroinflammatory Response in Microglial Cells Promotes Leukocyte Infiltration Across the BBB. Neurochem Res 41:193–209.

Alfonso-Loeches S, Ureña-Peralta JR, Morillo-Bargues MJ, Oliver-De La Cruz J, Guerri C. 2014. Role of mitochondria ROS generation in ethanol-induced NLRP3 inflammasome activation and cell death in astroglial cells. Front Cell Neurosci 8:216.

Allen-Worthington KH, Brice AK, Marx JO, Hankenson FC. 2015. Intraperitoneal Injection of Ethanol for the Euthanasia of Laboratory Mice (Mus musculus) and Rats (Rattus norvegicus). J Am Assoc Lab Anim Sci JAALAS 54:769–778.

Almeida-da-Silva CLC, Savio LEB, Coutinho-Silva R, Ojcius DM. 2023. The role of NOD-like receptors in innate immunity. Front Immunol 14:1122586.

Bang OY, Kim J-E. 2022. Stem cell-derived extracellular vesicle therapy for acute brain insults and neurodegenerative diseases. BMB Rep 55:20–29.

Brust V, Schindler PM, Lewejohann L. 2015. Lifetime development of behavioural phenotype in the house mouse (Mus musculus). Front Zool 12:S17.

Ergin K, Çetinkaya R. 2022. Regulation of MicroRNAs. Methods Mol Biol Clifton NJ 2257:1–32.

Fernandez-Lizarbe S, Pascual M, Guerri C. 2009. Critical role of TLR4 response in the activation of microglia induced by ethanol. J Immunol Baltim Md 1950 183:4733–4744.

Gandhi RM, Kogan CS, Messier C, Macleod LS. 2014. Visual-spatial learning impairments are associated with hippocampal PSD-95 protein dysregulation in a mouse model of fragile X syndrome. Neuroreport 25:255–261.

Hade MD, Suire CN, Suo Z. 2021. Mesenchymal Stem Cell-Derived Exosomes: Applications in Regenerative Medicine. Cells 10:1959.

Harris JA, Mihalas S, Hirokawa KE, Whitesell JD, Choi H, Bernard A, Bohn P, Caldejon S, Casal L, Cho A, Feiner A, Feng D, Gaudreault N, Gerfen CR, Graddis N, Groblewski PA, Henry AM, Ho A, Howard R, Knox JE, Kuan L, Kuang X, Lecoq J, Lesnar P, Li Y, Luviano J, McConoughey S, Mortrud MT, Naeemi M, Ng L, Oh SW, Ouellette B, Shen E, Sorensen SA, Wakeman W, Wang Q, Wang Y, Williford A, Phillips JW, Jones AR, Koch C, Zeng H. 2019. Hierarchical organization of cortical and thalamic connectivity. Nature 575:195– 202.

Hou B, Zhang Y, Liang P, He Y, Peng B, Liu W, Han S, Yin J, He X. 2020. Inhibition of the NLRP3-inflammasome prevents cognitive deficits in experimental autoimmune encephalomyelitis mice via the alteration of astrocyte phenotype. Cell Death Dis 11:377.

Hou L, Zhu Z, Jiang F, Zhao J, Jia Q, Jiang Q, Wang H, Xue W, Wang Y, Tian L. 2023. Human umbilical cord mesenchymal stem cell-derived extracellular vesicles alleviated silica induced lung inflammation and fibrosis in mice via circPWWP2A/miR-223-3p/NLRP3 axis. Ecotoxicol Environ Saf 251:114537.

Ibáñez F, Montesinos J, Ureña-Peralta JR, Guerri C, Pascual M. 2019. TLR4 participates in the transmission of ethanol-induced neuroinflammation via astrocyte-derived extracellular vesicles. J Neuroinflammation 16:136.

Jaszczyk A, Stankiewicz AM, Juszczak GR. 2022. Dissection of Mouse Hippocampus with Its Dorsal, Intermediate and Ventral Subdivisions Combined with Molecular Validation. Brain Sci 12:799.

Jin X, Liu M-Y, Zhang D-F, Zhong X, Du K, Qian P, Yao W-F, Gao H, Wei M-J. 2019. Baicalin mitigates cognitive impairment and protects neurons from microglia-mediated neuroinflammation via suppressing NLRP3 inflammasomes and TLR4/NF-κB signaling pathway. CNS Neurosci Ther 25:575–590.

Kanehisa M, Goto S. 2000. KEGG: kyoto encyclopedia of genes and genomes. Nucleic Acids Res 28:27–30.

Kelley N, Jeltema D, Duan Y, He Y. 2019. The NLRP3 Inflammasome: An Overview of Mechanisms of Activation and Regulation. Int J Mol Sci 20:3328.

Le Merre P, Ährlund-Richter S, Carlén M. 2021. The mouse prefrontal cortex: Unity in diversity. Neuron 109:1925–1944.

McClain JA, Hayes DM, Morris SA, Nixon K. 2011. Adolescent binge alcohol exposure alters hippocampal progenitor cell proliferation in rats: effects on cell cycle kinetics. J Comp Neurol 519:2697–2710.

Mellado S, Cuesta CM, Montagud S, Rodríguez-Arias M, Moreno-Manzano V, Guerri C, Pascual M. 2023. Therapeutic role of mesenchymal stem cell-derived extracellular vesicles in neuroinflammation and cognitive dysfunctions induced by binge-like ethanol treatment in adolescent mice. CNS Neurosci Ther.

Mellado-López M, Griffeth RJ, Meseguer-Ripolles J, Cugat R, García M, Moreno-Manzano V. 2017. Plasma Rich in Growth Factors Induces Cell Proliferation, Migration, Differentiation, and Cell Survival of Adipose-Derived Stem Cells. Stem Cells Int 2017:5946527.

Mira RG, Lira M, Tapia-Rojas C, Rebolledo DL, Quintanilla RA, Cerpa W. 2019. Effect of Alcohol on Hippocampal-Dependent Plasticity and Behavior: Role of Glutamatergic Synaptic Transmission. Front Behav Neurosci 13:288.

Montesinos J, Pascual M, Pla A, Maldonado C, Rodríguez-Arias M, Miñarro J, Guerri C. 2015. TLR4 elimination prevents synaptic and myelin alterations and long-term cognitive dysfunctions in adolescent mice with intermittent ethanol treatment. Brain Behav Immun 45:233–244.

Muñoz-Criado I, Meseguer-Ripolles J, Mellado-López M, Alastrue-Agudo A, Griffeth RJ, Forteza-Vila J, Cugat R, García M, Moreno-Manzano V. 2017. Human Suprapatellar Fat Pad-Derived Mesenchymal Stem Cells Induce Chondrogenesis and Cartilage Repair in a Model of Severe Osteoarthritis. Stem Cells Int 2017:4758930.

Nagel BJ, Schweinsburg AD, Phan V, Tapert SF. 2005. Reduced hippocampal volume among adolescents with alcohol use disorders without psychiatric comorbidity. Psychiatry Res 139:181–190.

Noori L, Arabzadeh S, Mohamadi Y, Mojaverrostami S, Mokhtari T, Akbari M, Hassanzadeh G. 2021. Intrathecal administration of the extracellular vesicles derived from human Wharton’s jelly stem cells inhibit inflammation and attenuate the activity of inflammasome complexes after spinal cord injury in rats. Neurosci Res 170:87–98.

Pan L, Yan B, Zhang J, Zhao P, Jing Y, Yu J, Hui J, Lu Q. 2022. Mesenchymal stem cells-derived extracellular vesicles-shuttled microRNA-223-3p suppress lipopolysaccharide-induced cardiac inflammation, pyroptosis, and dysfunction. Int Immunopharmacol 110:108910.

Pascual M, Blanco AM, Cauli O, Miñarro J, Guerri C. 2007. Intermittent ethanol exposure induces inflammatory brain damage and causes long-term behavioural alterations in adolescent rats. Eur J Neurosci 25:541–550.

Pascual M, López-Hidalgo R, Montagud-Romero S, Ureña-Peralta JR, Rodríguez-Arias M, Guerri C. 2021. Role of mTOR[regulated autophagy in spine pruning defects and memory impairments induced by binge[like ethanol treatment in adolescent mice. Brain Pathol 31:174–188.

Piancone F, La Rosa F, Marventano I, Saresella M, Clerici M. 2021. The Role of the Inflammasome in Neurodegenerative Diseases. Mol Basel Switz 26:953.

Qiu G, Zheng G, Ge M, Wang J, Huang R, Shu Q, Xu J. 2018. Mesenchymal stem cell-derived extracellular vesicles affect disease outcomes via transfer of microRNAs. Stem Cell Res Ther 9:320.

Quigley J, Committee on substance use and prevention. 2019. Alcohol Use by Youth. Pediatrics 144:e20191356.

Ru Y, Kechris KJ, Tabakoff B, Hoffman P, Radcliffe RA, Bowler R, Mahaffey S, Rossi S, Calin GA, Bemis L, Theodorescu D. 2014. The multiMiR R package and database: integration of microRNA-target interactions along with their disease and drug associations. Nucleic Acids Res 42:e133.

Saïd-Sadier N, Ojcius DM. 2012. Alarmins, inflammasomes and immunity. Biomed J 35:437–449.

Schmittgen TD, Livak KJ. 2008. Analyzing real-time PCR data by the comparative C(T) method. Nat Protoc 3:1101–1108.

Sultan FA. 2013. Dissection of Different Areas from Mouse Hippocampus. Bio-Protoc 3:e955.

Szklarczyk D, Kirsch R, Koutrouli M, Nastou K, Mehryary F, Hachilif R, Gable AL, Fang T, Doncheva NT, Pyysalo S, Bork P, Jensen LJ, von Mering C. 2023. The STRING database in 2023: protein-protein association networks and functional enrichment analyses for any sequenced genome of interest. Nucleic Acids Res 51:D638–D646.

Ureña-Peralta JR, Alfonso-Loeches S, Cuesta-Diaz CM, García-García F, Guerri C. 2018. Deep sequencing and miRNA profiles in alcohol-induced neuroinflammation and the TLR4 response in mice cerebral cortex. Sci Rep 8:15913.

Wang L, Hauenstein AV. 2020. The NLRP3 inflammasome: Mechanism of action, role in disease and therapies. Mol Aspects Med 76:100889.

Wu T, Hu E, Xu S, Chen M, Guo P, Dai Z, Feng T, Zhou L, Tang W, Zhan L, Fu X, Liu S, Bo X, Yu G. 2021. clusterProfiler 4.0: A universal enrichment tool for interpreting omics data. Innov Camb Mass 2:100141.

Xu F, Wu Y, Yang Q, Cheng Y, Xu J, Zhang Y, Dai H, Wang B, Ma Q, Chen Y, Lin F, Wang C. 2022. Engineered Extracellular Vesicles with SHP2 High Expression Promote Mitophagy for Alzheimer’s Disease Treatment. Adv Mater Deerfield Beach Fla 34:e2207107.

Yin K, Wang S, Zhao RC. 2019. Exosomes from mesenchymal stem/stromal cells: a new therapeutic paradigm. Biomark Res 7:8.

Zheng D, Liwinski T, Elinav E. 2020. Inflammasome activation and regulation: toward a better understanding of complex mechanisms. Cell Discov 6:36.

